# A reevaluation of the electrophysiological correlates of absolute pitch and relative pitch: No evidence for an absolute pitch-specific negativity

**DOI:** 10.1101/489302

**Authors:** Simon Leipold, Chantal Oderbolz, Marielle Greber, Lutz Jäncke

## Abstract

Musicians with absolute pitch effortlessly identify the pitch of a sound without an external reference. Previous neuroscientific studies on absolute pitch have typically had small samples sizes and low statistical power, making them susceptible for false positive findings. In a seminal study, Itoh et al. (2005) reported the elicitation of an absolute pitch-specific event-related potential component during tone listening — the *AP negativity*. Additionally, they identified several components as correlates of relative pitch, the ability to identify relations between pitches. Here, we attempted to replicate the main findings of Itoh et al.’s study in a large sample of musicians (n = 104) using both frequentist and Bayesian inference. We were not able to replicate the presence of an AP negativity during tone listening in individuals with high levels of absolute pitch, but we partially replicated the findings concerning the correlates of relative pitch. Our results are consistent with several previous studies reporting an absence of differences between musicians with and without absolute pitch in early auditory evoked potential components. We conclude that replication studies form a crucial part in assessing extraordinary findings, even more so in small fields where a single finding can have a large impact on further research.

## Introduction

A core pillar of scientific progress is the validation of the veracity of findings by means of replication studies. If an effect is true, it should be reliably obtained in an independent, adequately powered study using similar procedures (Simons, 2014). In recent years, the replicability of published findings has been repeatedly called into question. Meta-research studies have demonstrated, both theoretically and empirically, low replicability of findings across a diverse range of research fields (Baker, 2016; Begley and Ellis, 2012; Camerer et al., 2018, 2016; Ioannidis, 2005; Open Science Collaboration, 2015). Several reasons contributing to the low replicability of findings have been identified (Munafò et al., 2017). These include, for example, publication bias against negative results (Nissen et al., 2016; Rosenthal, 1979), questionable research practices (Kerr, 1998; Neuroskeptic, 2012; Simmons et al., 2011), misunderstandings of p values (Halsey et al., 2015), and maybe most importantly, low statistical power (Button et al., 2013).

Not only does low statistical power result in a decreased probability of finding an effect should a true effect exist, it also reduces the probability of a significant effect being a true effect (Button et al., 2013). Cognitive neuroscience represents a research field wherein the problem of low statistical power is especially evident (Nord et al., 2017; Szucs and Ioannidis, 2017). The use of non-invasive techniques such as functional magnetic resonance imaging (fMRI) and electroencephalography (EEG) on human subjects is both time-consuming and resource-intensive; thus, cognitive neuroscience studies often have small sample sizes which are only adequate to detect very large effects (Poldrack et al., 2017). In addition, the problem of small samples is aggravated in studies investigating rare populations with unique characteristics because the subject pool of these studies is inherently limited.

A prime example for studies on rare populations using small samples sizes are neuroscientific studies on individuals with absolute pitch (AP) — the ability to effortlessly name the pitch of a sound without an external reference sound (Deutsch, 2013). In the last 25 years, these studies have provided important insights into the cognitive, neuroanatomical, and neurophysiological foundations of this ability (Keenan et al., 2001; Schlaug et al., 1995; Zatorre et al., 1998). However, many of the findings have not yet been replicated. In 2005, a seminal EEG study on AP was published in *Cerebral Cortex* (Itoh et al., 2005). The authors evaluated four groups of subjects with varying levels of AP (High-AP musicians, Mid-AP musicians, Low-AP musicians, and untrained individuals without AP). Each of the groups consisted of 11 subjects. The main finding of Itoh et al.’s study was the unique elicitation of a left posterior temporal negativity in High-AP musicians in response to pure tones. This component of the event-related potential (ERP) occurring 150 ms after stimulus-onset was termed *AP negativity* because it presumably reflected the automatic retrieval of the association of a pitch and its label (e.g., C#) in High-AP musicians. Further findings of Itoh et al.’s study concerned the ERP correlates of relative pitch (RP), the ability to identify the relation between successive pitches either by making higher-lower judgments or by determining the exact musical interval between the pitches (McDermott and Oxenham, 2008). The authors found three ERP components (P3b, parietal positive slow wave, frontal negative slow wave) occurring later than 300 ms after stimulus-onset that were not elicited in High-AP musicians and gradually increased with lower levels of AP. Thus, these components presumably reflected the cortical processing related to RP.

In small research fields such as those exploring AP, single studies with extraordinary findings can have a large impact on subsequently conducted studies. Itoh et al.’s study, a prime example of such a case, was the first to find an electrophysiological marker of AP occurring as early as 150 ms after stimulus-onset during simple tone listening. Subsequent studies have cited this aspect of the study as part of the motivation for their use of EEG to investigate the timing of the neurophysiological responses in AP (e.g., Elmer et al., 2013; Rogenmoser et al., 2015). Furthermore, many studies cite that the AP negativity was (exclusively) found at an electrode over the left temporal cortex as evidence for the importance of the left-sided planum temporale in AP (Loui et al., 2012; Oechslin et al., 2010; e.g., Schulze et al., 2009). Surprisingly, since the publication of Itoh et al.’s study more than 10 years ago, no study has been able to replicate the finding of an AP negativity in musicians with high levels of AP. This is in stark contrast to the findings concerning the ERP correlates of RP, which are consistent with numerous studies finding an absent or reduced P3b component in AP musicians (Hantz et al., 1992; Klein et al., 1984; Wayman et al., 1992).

In several recent high-profile publications, the need for replications and its scientific value has been strongly emphasized (Munafò et al., 2017; Poldrack et al., 2017; Zwaan et al., 2018). There have been first attempts to replicate findings in cognitive neuroscience (e.g., Boekel et al., 2015; Nieuwland et al., 2018). In light of these developments, we attempted to replicate the main findings of the study by Itoh et al. using closely matched materials, procedures, and analyses. We recruited a sample of 104 musicians — more than double the sample size of the original study. These subjects performed a pitch-naming task during EEG acquisition. We evaluated the AP negativity as well as the ERP components related to RP, namely the P3b, the parietal positive slow wave (ppSW), and the frontal negative slow wave (fnSW). To quantify the success of the replication, we used Bayesian inference along with the frequentist inference employed in the original study.

## Materials and Methods

### Subjects

We examined 104 musicians with varying degrees of AP that were assigned to three groups (High-AP musicians, Mid-AP musicians, Low-AP musicians) according to their tone-naming proficiency (see Figure 1). The group assignment was based on two cutoffs in tone-naming scores (37.5%, 82.5%) that were chosen to optimize the balance between, on the one hand, a close matching of the group-specific tone-naming scores in the original study and, on the other hand, a high enough number of subjects in each group. Note that the cutoff scores chosen are specific to the sample investigated in this study and should not be taken as non-arbitrary benchmarks for future studies. In the original study, the criterion for group assignment was not reported. Unlike in the original study, we did not include a group without musical training for two reasons: It would have been difficult for them to understand the instructions of the EEG task, and their results would have been hard to interpret because of the confounding factor of musical training.

**Figure 1.**
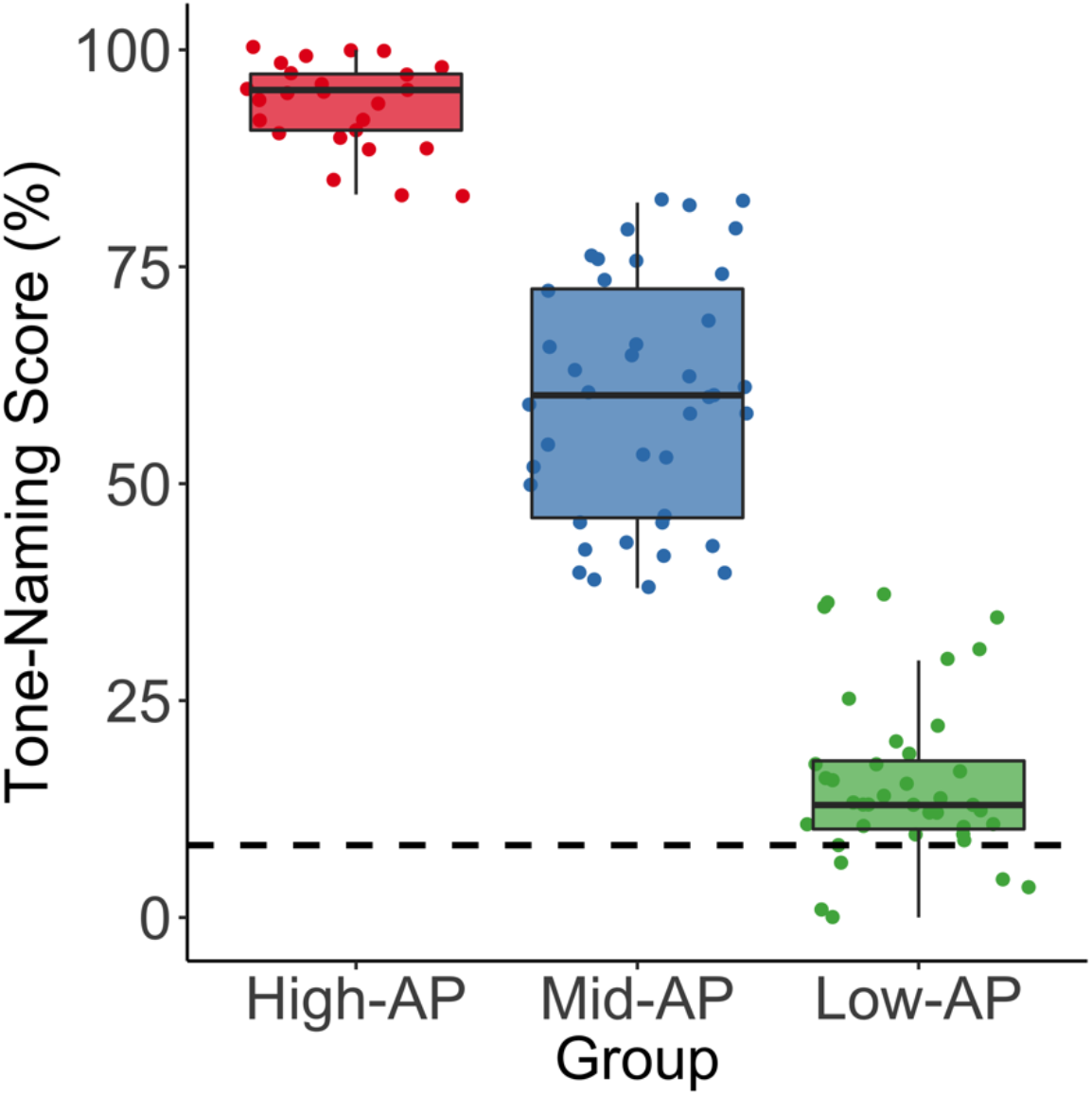
Distribution of tone-naming scores across groups. The subjects (n = 104) were assigned to three groups — High-AP musicians, Mid-AP musicians, and Low-AP musicians — according to their tone-naming proficiency. The assignment was based on two cutoffs in tone-naming scores (37.5%, 82.5%). These values were chosen to optimize the balance between a close matching of the group-specific tone-naming scores in the original study and a sufficient number of subjects in each group.

All subjects were either professional musicians, music students, or highly trained amateurs between 18 and 44 years of age, who were recruited in the context of a larger project investigating AP. None of the subjects reported any neurological, audiological, or severe psychiatric disorders. Pure tone audiometry (ST20, MAICO Diagnostics, Berlin, Germany) confirmed the absence of hearing loss in all subjects. The demographical data (sex, age, handedness) and part of the behavioral data (tone-naming proficiency and musical experience) were collected using an online survey tool (http://www.limesurvey.org/). Self-reported handedness was verified using a German translation of the Annett questionnaire (Annett, 1970). Subjects provided written informed consent and were paid for their participation. The study was approved by the local ethics committee (http://www.kek.zh.ch/) and conducted according to the principles defined in the Declaration of Helsinki.

### Tone-Naming Test

To assess their tone-naming ability, all subjects completed a tone-naming test (Oechslin et al., 2010). During the test, the subjects had to name both the chroma and the octave (e.g. E4) of 108 pure tones which were presented in a pseudorandomized order. The tones had a duration of 500 ms and were masked with 2000 ms Brownian noise presented immediately before and after each tone. The test included all tones from C3 to B5 (twelve-tone equal temperament tuning, A4 = 440 Hz). Each of the tones was presented three times. The tone-naming score was calculated using the percentage of correct chroma identifications without considering octave errors (Deutsch, 2013). Therefore, the chance-level was at 8.3%.

### Experimental Procedure

During EEG data acquisition, the subjects performed a slightly modified version of the pitch-naming task employed in the original study. Itoh et al. also included a Stroop-like ERP experiment in their study which we chose not to replicate because in the original study, it mainly served as an attempted internal replication of the identified ERP correlates of AP and RP. Thus, in this study, we do not discuss the findings of this Stroop-like experiment.

All stimuli and the stimulus presentation scripts are available online (https://osf.io/93gdm/). We used identical auditory stimuli to the ones used in the original study. The stimuli consisted of three pure tones with different frequencies (262 Hz, 294 Hz, and 330 Hz), corresponding to C4, D4, and E4 in twelve-tone equal temperament tuning. Note that in Itoh et al.’s publication, the tones were labeled incorrectly (C3, D3, and E3). All tones had the same temporal envelope, characterized by a duration of 350 ms, a 10 ms linear fade-in, and a 50 ms linear fade-out. The stimuli were created using Audacity (version 2.1.2, http://www.audacityteam.org/) and presented at a sound pressure level of 75 dB via on-ear headphones (HD 25-1, Sennheiser, Wedemark, Germany) using Presentation software (version 18.1, www.neurobs.com).

The pitch-naming task consisted of a Listening and a Labeling condition (the Naming condition of the original study). Both experimental conditions encompassed 180 trials, and each of the three pure tones was presented 60 times. The trials were presented in a randomized order. Within a single trial, first, a pure tone was presented, followed by a silent inter-stimulus interval (jittered duration = 900 ms – 1100 ms) after which an auditory cue (pink noise, duration = 10 ms, linear fade-in = 2 ms, linear fade-out = 2 ms) was presented. This cue indicated to the subjects that they should respond by a key press. The trial continued in silence until a response was given. After the response, there was an inter-trial interval (duration = 1000 ms), before the next pure tone was presented. The experimental conditions only differed in terms of the instructions given to the subjects: In the Listening condition, the subjects listened to the tones and pressed a neutrally marked key in response to the auditory cue, irrespective of the chroma of the pure tone that was presented before. In the Labeling condition, the subjects labeled the pure tones by pressing one of three corresponding keys that were marked with the tone names (C, D, and E), also in response to the auditory cue. In both conditions, the subjects were instructed to respond as quickly and as accurately as possible. Importantly, no responses were given directly after the tone in order to avoid a contamination of the ERPs by motor artifacts. Before each condition, we included six practice trials to familiarize the subjects with the task. During both experimental conditions, a screen in front of the subjects showed a black fixation cross on a gray background. The whole task lasted around 20 minutes.

Our version of the pitch-naming task differed from Itoh et al.’s version in three ways: First, we doubled the number of trials per condition (180 instead of 90) to increase the signal to noise ratio and therefore the statistical power. Second, in order to accurately record the subjects’ responses and to avoid articulation-related artifacts, we chose to refrain from using verbal responses. In the original study, the subjects responded verbally. Third, we fixed the order of the conditions across all subjects whereas in the original study, the order was balanced across subjects. The rationale for this decision is as follows: Group differences in ERP amplitudes during Listening are interpreted in relation to the AP-specific automaticity of the retrieval of the pitch-label association that occurs even without external instruction. Consequently, in order to avoid the contamination of the Listening condition by possible spill-overs from the instruction of a preceding Labeling condition, we decided to always present the Listening condition before the Labeling condition.

### EEG Data Acquisition and Preprocessing

The EEG data was acquired continuously using an electrode cap (Easycap, Herrsching, Germany) with 32 Ag / AgCl electrodes placed according to an extended 10/20 system (Fp1, Fp2, F7, F3, Fz, F4, F8, FT7, FC3, FCz, FC4, FT8, T7, C3, Cz, C4, T8, TP9, TP7, CP3, CPz, CP4, TP8, TP10, P7, P3, Pz, P4, P8, O1, Oz, O2) and a BrainAmp amplifier (Brainproducts, Munich, Germany). The reference electrode was placed on the tip of the nose. We used a sampling rate of 1000 Hz and an online bandpass filter between 0.1 Hz and 100 Hz. Electrode impedance was kept below 10 kΩ throughout the acquisition using electrically conductive gel.

The preprocessing of the EEG data was performed using BrainVision Analyzer (Version 2.1, https://www.brainproducts.com/). We bandpass-filtered the data from 0.5 Hz to 20 Hz (48 dB/octave) and applied a notch filter of 50 Hz. Artifacts caused by eye blinks and saccades were corrected using independent component analysis (Jung et al., 2000) with the remaining artifacts being removed using an automatic raw data inspection (removal criteria: amplitude gradient > 50 µV/ms, amplitude difference > 100 µV, amplitude minimum/maximum > −100 µV / 100 µV). Afterwards, we segmented the continuous data into epochs of 900 ms (−100 ms to 800 ms relative to pure-tone onset). Finally, we baseline-corrected these epochs using the interval from −100 ms to the onsets of the pure tones.

The baseline-corrected epochs were averaged per subject and experimental condition to compute ERPs. From these ERPs, we extracted the mean amplitudes of the following time intervals, originally specified in Itoh et al.’s study: P3b = 300 – 450 ms, ppSW = 450 – 550 ms, fnSW = 550 ms – 800 ms, AP negativity = 145 – 155 ms. Note that for the fnSW, Itoh et al. used an interval from 550 ms to 900 ms (the end of their epochs). In our case, however, due to the jittered inter-stimulus interval, some of the auditory cues already appeared 900 ms after the onset of the pure tones and thus, to avoid motor artifacts from key presses in anticipation of the auditory cue, we reduced the designated interval by 100 ms. The extracted mean amplitudes were then subjected to statistical analysis. We only analyzed 20 of the 32 electrodes to match the electrode locations used in the original study. The analyzed electrodes were as follows: Fp1, Fp2, F7, F3, Fz, F4, F8, T7, C3, Cz, C4, T8, P7, P3, Pz, P4, P8, O1, Oz, O2. Note that in the original study, the electrode locations T7, T8, P7, and P8 are called T3, T4, T5, and T6 respectively, in line with the classical 10-20 system terminology. Thus, in the following, we refer to these electrode locations as T7/T3, T8/T4, P7/T5 and P8/T6.

### Statistical analysis

The statistical analysis of the EEG data and the behavioral data was performed in R (version 3.3.2, http://www.r-project.org/). In addition to the frequentist inference employed in the original study, we used Bayesian inference, more specifically Bayes factors, to quantify the evidence for the alternative hypothesis relative to the null hypothesis and vice versa (Kass and Raftery, 1995). In contrast to frequentist inference, which can only be used to reject the null hypothesis, Bayesian inference allows statements concerning the evidence in support of the null hypothesis (Dienes, 2011). As such, Bayesian inference is ideally suited for replication studies to avoid the non-interpretability of a non-significant effect obtained using frequentist inference (Anderson and Maxwell, 2016). Consequently, in addition to p values, we report Bayes factors favoring either the null hypothesis (BF_01_) or the alternative hypothesis (BF_10_). Bayes factors are interpreted in a straightforward way: A BF_10_ of 3 indicates that the observed data is three times more likely under the alternative hypothesis than under the null hypothesis. In this study, a Bayes factor between 1 and 3 is considered as anecdotal evidence, a Bayes factor between 3 and 10 as moderate evidence, and a Bayes factor between 10 and 30 as strong evidence for one hypothesis relative to the other hypothesis (Boekel et al., 2015; Jeffreys, 1961). To calculate Bayesian t-tests (Rouder et al., 2009) and Bayesian ANOVAs (Rouder et al., 2012), we used the R package *BayesFactor* (version 0.9.12-2, https://CRAN.R-project.org/package=BayesFactor). We used the default priors as implemented in the *BayesFactor* package (scale value *r* = 0.707). The use of default priors is advantageous because these priors do not depend on effect size estimates drawn from previous studies which are often known to be inflated, especially in studies with small sample sizes (Ioannidis, 2008). Nonetheless, we checked a range of different scale values, which did not change the conclusions drawn from the resulting Bayes factors. Hence, only Bayes factors based on the default scale value are reported. To calculate frequentist ANOVAs, we used the R package *ez* (version 4.4.0, https://CRAN.R-project.org/package=ez). In case of non-sphericity, Greenhouse-Geisser corrected degrees of freedom and p values are reported. The significance level was set to α = 0.05 for all analyses unless otherwise stated. Effect sizes within an ANOVA are given as generalized eta-squared (η^2^_G_) and effect sizes for t-tests are given as Cohen’s d (*d*).

### Behavioral Data Analysis

The behavioral and demographical subject characteristics were compared between the three groups using a one-way ANOVA per characteristic. The behavioral measures acquired during the EEG experiment (response accuracy and response time) were analyzed separately. For the response time, we calculated a two-way mixed-design ANOVA with a within-subject factor Condition and a between-subject factor Group. Trials with response times shorter than 200 ms or longer than 1000 ms were excluded from the analysis. For the response accuracy in the Labeling condition, we calculated a one-way ANOVA with a between-subject factor Group and subsequently performed pairwise comparisons (Bonferroni adjusted α = 0.017). In the Listening condition, there was no response choice involved and thus, no response accuracy was calculated.

### EEG Data Analysis

For each of the ERP components related to RP processing — P3b, ppSW, and fnSW — we first calculated a two-way mixed-design ANOVA with a within-subject factor Condition (Listening vs. Labeling) and a between-subject factor Group (High-AP vs. Mid-AP vs. Low-AP) at the electrode locations identified in the original study as showing a statistically significant Group x Condition interaction. Consequently, we analyzed the amplitudes of both the P3b and the ppSW at electrode Pz. As we did not collect data for electrode Fpz, we calculated and analyzed the mean amplitudes of electrodes Fp1 and Fp2 for the fnSW. In case of a statistically significant Group x Condition interaction, for each group we calculated the difference in amplitudes between the conditions (Labeling minus Listening). These differences were then compared pairwise between the groups using two sample t-tests (Bonferroni adjusted α = 0.017). Note that in the original study, differential condition effects between the groups were inferred based on a statistically significant condition effect in one group and a simultaneous non-significant condition effect in another group. Although somewhat common in earlier neuroscientific research, this procedure is erroneous in that a comparison of condition effects requires a statistical test on their difference (Gelman and Stern, 2006; Nieuwenhuis et al., 2011). Nevertheless, to ensure comparability with the original study, we additionally implemented this procedure in order to check whether a component was elicited in one group, but not in another. This was done by comparing the amplitudes of the two conditions within each group separately using a paired t-test (Bonferroni adjusted α = 0.017).

To explore potential effects at unexpected electrodes and to ensure comparability with the original study, for each RP-related component, we additionally calculated a three-way mixed-design ANOVA with the two within-subject factors Condition and Electrode (all 20 electrodes) and a between-subject factor Group. We did not calculate Bayes factors for the three-way ANOVAs because they are not well-suited for designs of high complexity (Rouder et al., 2017). In case of a three-way interaction, a two-way ANOVA with the factors Condition and Group was performed at each of the 20 electrodes (Bonferroni adjusted α = 0.0025). We used the conservative Bonferroni correction because in an N-way ANOVA with a large number of factor levels, the family-wise error (FWE) rate is already inflated at the level of the ANOVA itself (Cramer et al., 2016; Luck and Gaspelin, 2017). Therefore, in follow-up ANOVAs, it is crucial to adequately correct for the inflated FWE rate. For electrodes showing a statistically significant Group x Condition interaction, we again calculated condition differences and compared these differences pairwise between the groups using two sample t-tests (Bonferroni adjusted α = 0.017). Finally, we again checked whether a component was elicited in one group, but not another. It is important to note that it is generally not advisable to compute N-way ANOVAs with a large number of factor levels as it was done in the original study. As described above, on the one hand, in these types of ANOVAs the FWE rate and thus the probability for a false-positive finding increases. On the other hand, an adequate adjustment of the significance level to correct for the inflated FWE rate decreases the statistical power of the study (Luck and Gaspelin, 2017).

The ERP component related to AP processing (AP negativity) was only analyzed in the Listening condition at electrode P7/T5 using a one-way ANOVA with a between-subject factor Group. In the original study, group differences were again erroneously inferred based on a significant effect in one group and a simultaneous non-significant effect in the other group (Gelman and Stern, 2006; Nieuwenhuis et al., 2011). To ensure comparability with the original study, we additionally checked in which of the groups the component was elicited above baseline, using a t-test against zero in each group (Bonferroni adjusted α = 0.017).

## Results

### Behavior

The three groups did not differ with regard to age (F(2,101) = 1.46, p = 0.24, BF_01_ = 3.28, η^2^_G_ = 0.03) and years of musical training (F(2,101) = 1.47, p = 0.24, BF_01_ = 3.31, η^2^_G_ = 0.03). As expected, the three groups substantially differed in their tone-naming proficiency (F(2,101) = 422.12, p < 10^-49^, BF_10_ > 10^46^, η^2^_G_ = 0.89). High-AP musicians had a higher tone-naming score than Mid-AP musicians (t(63) = 11.65, p < 0.001, BF_10_ > 10^14^, *d* = 2.97) and Mid-AP musicians had a higher score than Low-AP musicians (t(77) = 16.36, p < 0.001, BF_10_ > 10^23^, *d* = 3.68). Note that both in the original study and in this study, the subjects of the Low-AP group performed only slightly above chance-level. Therefore, these subjects represent what previous studies have termed Non-AP or RP musicians. To be consistent with the terminology of the original study, we are also using the term Low-AP musicians. Descriptive statistics of the subject characteristics are given in Table 1.

**Table 1:** Subject characteristics. Continuous measures given as mean ± standard deviation.

|  | High-AP musicians | Mid-AP musicians | Low-AP musicians |
| --- | --- | --- | --- |
| Number of subjects | 25 | 40 | 39 |
| Sex (female / male) | 12 / 13 | 19 / 21 | 20 / 19 |
| Handedness (right / left / both) | 23 / 1 / 1 | 34 / 4 / 2 | 35 / 3 / 1 |
| Age (years) | 25.80 ± 4.74 | 27.02 ± 5.72 | 25.10 ± 4.45 |
| Tone-naming score (%) | 93.59 ± 5.02 | 59.70 ± 13.96 | 15.69 ± 9.45 |
| Training (years) | 20.40 ± 4.54 | 20.62 ± 6.80 | 18.62 ± 4.69 |

Abbreviations: AP = absolute pitch

The two-way ANOVA of the response time revealed no main effect of Group (F(2,101) = 1.69, p = 0.19, BF_01_ = 1.66, η^2^_G_ = 0.03), no main effect of Condition (F(1,101) = 0.001, p = 0.98, BF_01_ = 6.77, η^2^_G_ < 0.001), and no Group x Condition interaction (F(2,101) = 0.46, p = 0.63, BF_01_ = 7.94, η^2^_G_ = 0.001). Because the subjects were instructed to press a key in response to a delayed cue and not in response to the tones itself, these response times should only be interpreted as markers of attentional or motivational processes and not with regard to the processing of the tones itself. Accordingly, there were no systematic differences between the groups or the conditions concerning the subjects’ attention or motivation during the experiment. The one-way ANOVA of the response accuracy in the Labeling condition revealed a main effect of Group (F(2,101) = 4.62, p = 0.01, BF_10_ = 3.90, η^2^_G_ = 0.08), in which Mid-AP musicians showed a higher response accuracy than Low-AP musicians (t(77) = 2.78, p = 0.007, BF_10_ = 6.16, *d* = 0.63). However, High-AP musicians did not show a higher response accuracy than Mid-AP musicians (t(63) = −0.99, p = 0.32, BF_01_ = 2.55, *d* = 0.25) and Low-AP musicians (t(62) = 1.63, p = 0.11, BF_01_ = 1.27, *d* = 0.42). Consistent with the response accuracies reported in the original study, all groups performed on a high level: mean (± standard deviation) of High-AP = 99.56% (± 0.89), Mid-AP = 99.74% (± 0.58), and Low-AP = 98.97% (± 1.63).

### Electrophysiological correlates of relative pitch

The group-averaged ERPs per condition at electrodes Fz, Cz, and Pz are shown in Figure 2. Condition differences are clearly visible in later ERP components (starting from 300 ms after pure tone onset). Group-averaged difference waveforms (Labeling minus Listening) and difference topographies for the P3b and the ppSW components are shown in Figure 3. In all groups, a P3b and a ppSW are clearly discernible from the difference waveforms. Both components were maximally elicited at parietal electrodes. Group-averaged difference waveforms for the fnSW component are shown in Figure 4A. The ERPs and topographies were visualized using functions from the R package *eegUtils* (Craddock, 2018).

**Figure 2.**
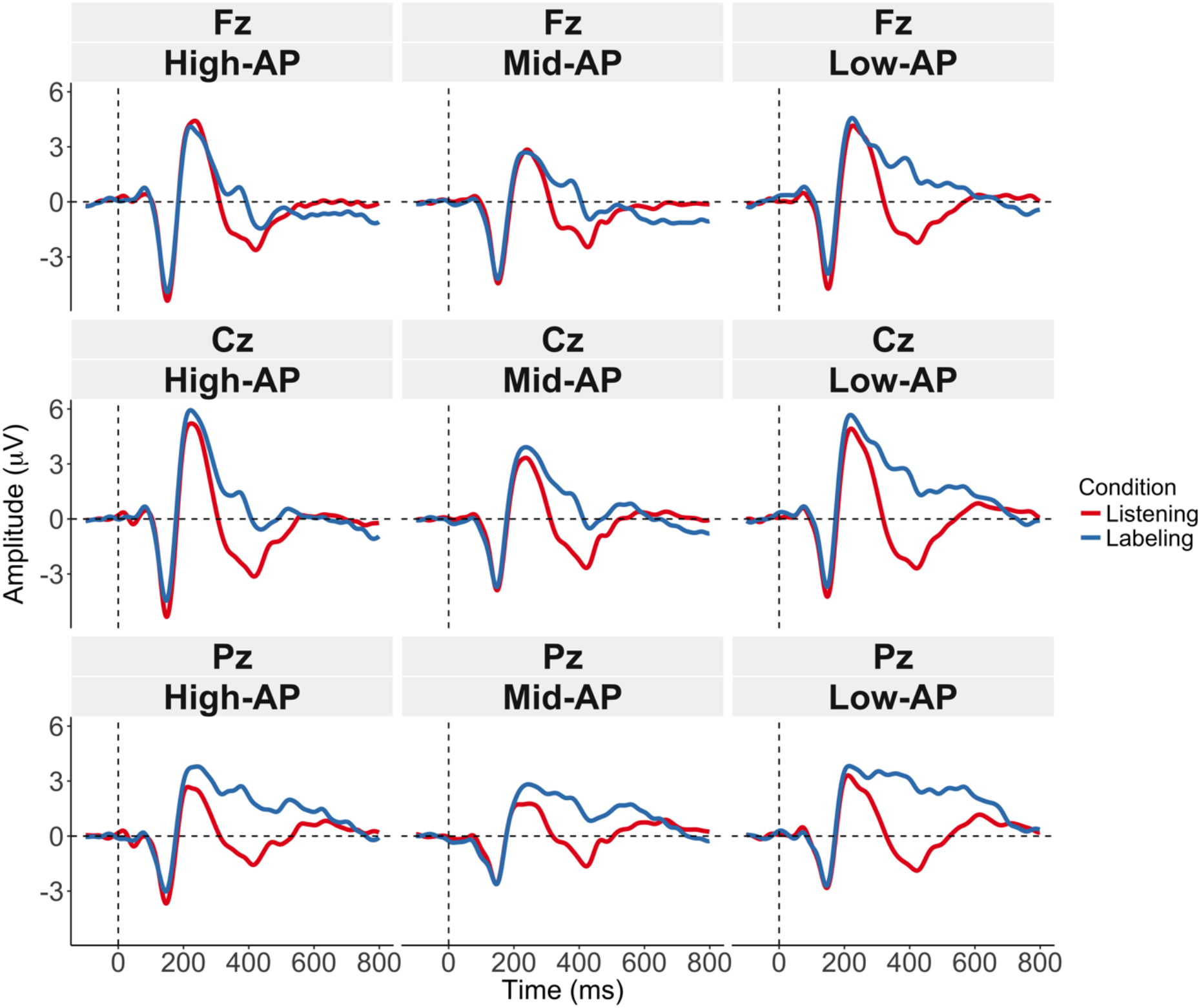
Event-related potential waveforms. Group-averaged ERP waveforms per experimental condition at electrodes Fz (upper panels), Cz (middle panels), and Pz (lower panels). The ERPs show the typical characteristics of an auditory evoked potential, including an N1-P2 complex. Condition differences are clearly visible in components elicited later than 300 ms after pure-tone onset.

**Figure 3.**
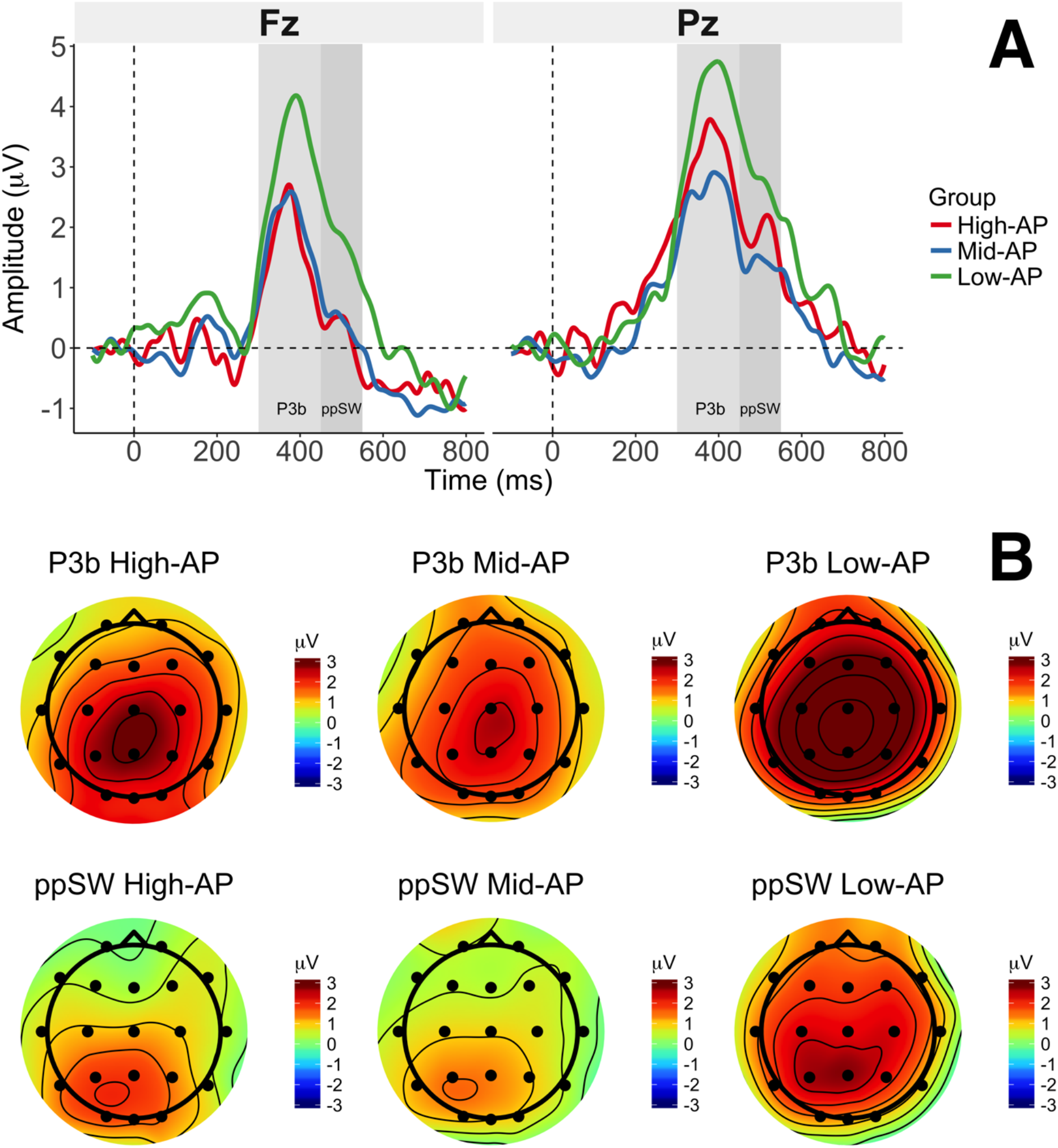
Replicated ERP correlates of relative pitch. Difference waveforms and topographies (Labeling minus Listening) showing the P3b and parietal positive slow wave (ppSW) components. (A) The group-specific difference waveforms (Labeling minus Listening) showed clearly discernible P3b (light gray-shaded) and ppSW (dark gray-shaded) components. At electrode Fz, Low-AP musicians showed a larger P3b than Mid-AP musicians and High-AP musicians. There were no differences between Mid-AP and High-AP musicians. The ppSW was not analyzed at Fz because the two-way interaction (Group x Condition) at this electrode did not reach statistical significance. At electrode Pz, Low-AP musicians showed a larger P3b than Mid-AP musicians, but we found no P3b differences between Low-AP and High-AP musicians and between Mid-AP and High-AP musicians. Furthermore, Low-AP musicians showed a larger ppSW than Mid-AP musicians, but there were no ppSW differences between Low-AP and High-AP musicians and between Mid-AP and High-AP musicians. (B) The group-specific difference topographies (Labeling minus Listening) showed a maximal elicitation of the P3b and the ppSW over parietal electrodes in all groups. Low-AP musicians showed a more extensive elicitation of both components.

#### P3b

The two-way ANOVA of the P3b amplitude at electrode Pz revealed a Group x Condition interaction (F(2,101) = 3.86, p = 0.02, BF_10_ = 2.31, η^2^_G_ = 0.02). The pairwise comparisons of condition differences in the P3b revealed that Low-AP musicians showed a larger difference than Mid-AP musicians (t(77) = 2.96, p = 0.004, BF_10_ = 9.38, *d* = 0.67). In contrast, there were no differences between Low-AP and High-AP musicians (t(62) = 1.32, p = 0.19, BF_01_ = 1.84, *d* = 0.34) and between Mid-AP and High-AP musicians (t(63) = −1.00, p = 0.32, BF_01_ = 2.53, *d* = 0.25). At electrode Pz, the P3b component was elicited in Low-AP musicians (t(38) = 9.46, p < 0.001, BF_10_ > 10^8^, *d* = 1.51), in Mid-AP musicians (t(39) = 8.33, p < 0.001, BF_10_ > 10^7^, *d* = 1.32), and in High-AP musicians (t(24) = 5.26, p < 0.001, BF_10_ = > 10^3^, *d* = 1.05).

The three-way ANOVA of the P3b amplitude using all electrodes revealed a Group x Condition x Electrode interaction (F(6.41,323.46) = 2.72, p = 0.01, η^2^_G_ = 0.002). The follow-up two-way ANOVAs revealed a Group x Condition interaction at electrode Fz (F(2,101) = 6.67, p = 0.002, BF_10_ = 18.81, η^2^_G_= 0.01). The pairwise comparison of condition differences revealed that Low-AP musicians showed a higher difference than Mid-AP musicians (t(77) = 3.13, p = 0.002, BF_10_ = 14.32, *d* = 0.71) and Low-AP musicians also showed a higher difference than High-AP musicians (t(62) = 2.98, p = 0.004, BF_10_ = 9.51, *d* = 0.76). However, there were no differences between Mid-AP musicians and High-AP musicians (t(63) = 0.52, p = 0.60, BF_01_ = 3.43, *d* = 0.13). At electrode Fz, again, the component was elicited in Low-AP musicians (t(38) = −10.21, p < 0.001, BF_10_ > 10^9^, *d* = 1.64), in Mid-AP musicians (t(39) = −7.32, p < 0.001, BF_10_ > 10^6^, *d* = 1.16), and in High-AP musicians (t(24) = −4.08, p < 0.001, BF_10_ = 73.29, *d* = 0.82).

#### Parietal Positive Slow Wave (ppSW)

The two-way ANOVA of the ppSW amplitude at electrode Pz revealed a Group x Condition interaction (F(2,101) = 3.30, p = 0.04, BF_10_ = 1.41, η^2^_G_ = 0.02). According to the pairwise comparisons of condition differences, Low-AP musicians showed a larger condition difference in ppSW amplitude than Mid-AP musicians (t(77) = 2.77, p = 0.007, BF_10_ = 6.04, *d* = 0.62). In contrast there were no differences between Low-AP and High-AP musicians (t(62) = 1.32, p = 0.19, BF_01_ = 1.85, *d* = 0.34) and between Mid-AP and High-AP musicians (t(63) = −0.80, p = 0.43, BF_01_ = 2.95, *d* = 0.20). The ppSW component was elicited in all groups: Low-AP (t(38) = 7.02, p < 0.001, BF_10_ > 10^5^, *d* = 1.12), Mid-AP (t(39) = 4.31, p < 0.001, BF_10_ = 229.35, *d* = 0.68), and High-AP (t(24) = 3.09, p = 0.005, BF_10_ = 8.64, *d* = 0.62). The three-way ANOVA of the ppSW amplitude using all electrodes revealed a Group x Condition x Electrode interaction (F(6.06,306.10) = 2.89, p = 0.01, η^2^_G_ = 0.004). However, the follow-up two-way ANOVAs did not reveal a Group x Condition at any electrode (all p > Bonferroni adjusted α).

#### Frontal Negative Slow Wave (fnSW)

The two-way ANOVA of the fnSW amplitude at electrode location Fcz (calculated as the mean of electrodes Fp1 and Fp2) did not reveal a Group x Condition interaction (F(2,101) = 0.92, p = 0.40, BF_01_ = 5.10, η^2^_G_ = 0.003). Furthermore, the three-way ANOVA using all electrodes did not reveal a Group x Condition x Electrode interaction (F(5.92,298.82) = 1.77, p = 0.11, η^2^_G_ = 0.002). Note that for this analysis, we only used 19 electrodes because the mean amplitude of electrodes Fp1 and Fp2 (corresponding to electrode Fcz) was used instead of these electrodes themselves.

### Electrophysiological correlates of absolute pitch

Figure 4b visualizes the group-averaged waveforms in the Listening condition at medial temporal and posterior temporal electrodes. Itoh et al. found a High-AP group-specific temporal negativity at left-sided electrode P7/T5 with a latency of 150 ms (AP negativity). In this study, the waveforms did not clearly show an elicitation of an AP negativity in any of the groups (see Figure 4b, bottom-left panel).

**Figure 4.**
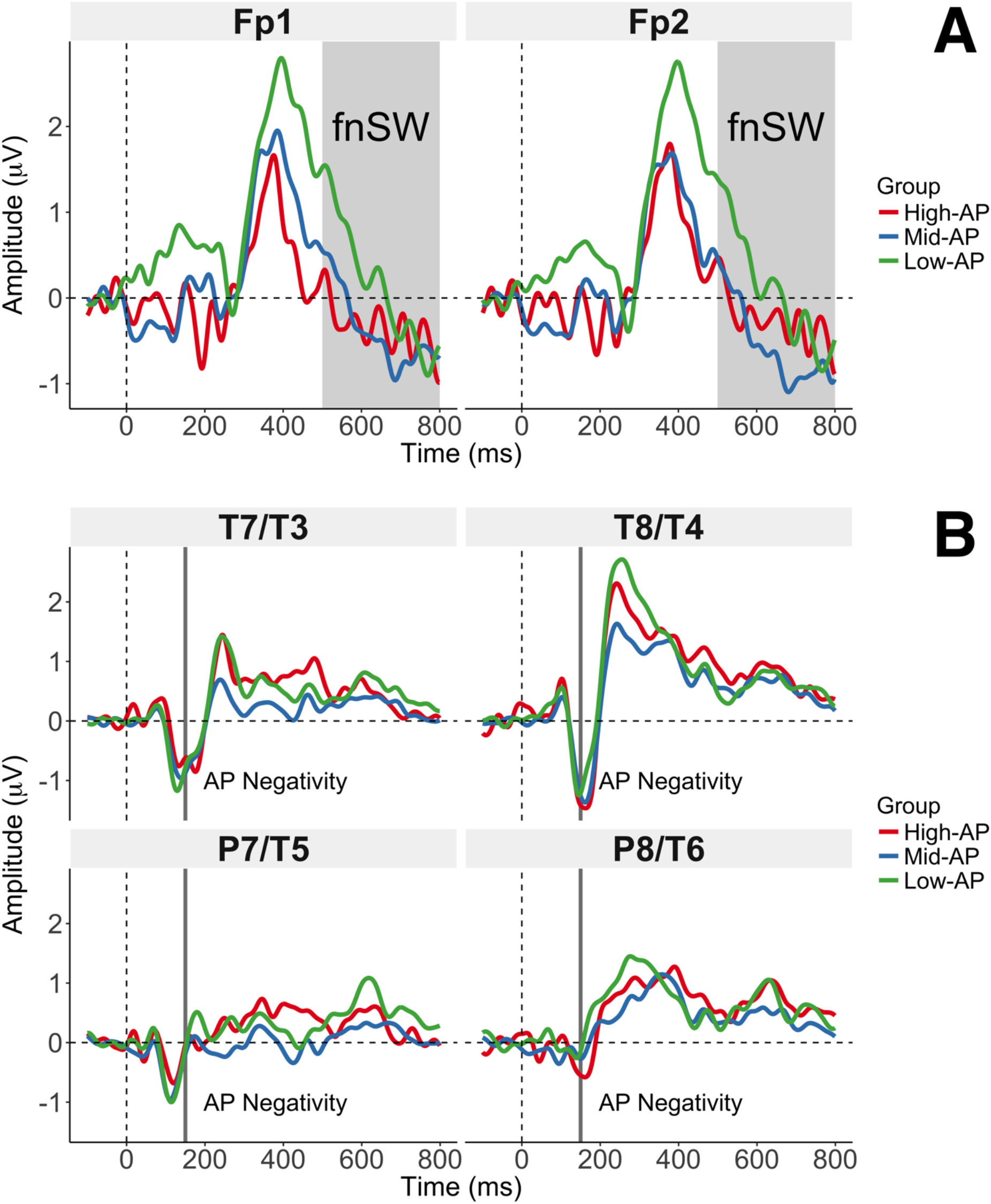
Non-replicated ERP correlates of relative and absolute pitch. (A) Group-averaged difference waveforms (Labeling minus Listening) at electrodes Fp1 and Fp2 showing the frontal negative slow wave (fnSW). In contrast to the original study, we found no group differences in the fnSW (gray-shaded area). Note that we analyzed the fnSW as the mean of Fp1 and Fp2 as we did not collect data at electrode location Fpz. (B) Group-averaged waveforms in the Listening condition at medial temporal (T7/T3, T8/T4) and posterior temporal electrodes (P7/T5, P8/T6). Contrary to the original study, we found no elicitation of an AP negativity in any of the groups at electrode P7/T5 (bottom-left panel; gray-shaded area). Consistent with the original study, there were no group differences in the other temporal electrodes (bottom-right panel and upper panels; gray-shaded areas).

#### Left Posterior Temporal Negativity (AP Negativity)

The one-way ANOVA of the AP negativity at electrode P7/T5 in the Listening condition did not reveal a main effect of Group (F(2,101) = 0.04, p = 0.96, BF_01_ = 10.61, η^2^_G_ < 0.001). As shown in Figure 5, the component was not elicited in any of the three groups: Low-AP (t(38) = −0.38, p = 0.71, BF_01_ = 5.42, *d* = 0.06), Mid-AP (t(39) = −0.63, p = 0.53, BF_01_ = 4.86, *d* = 0.09), and, most importantly, High-AP (t(24) = −0.11, p = 0.91, BF_01_ = 4.72, *d* = 0.02).

**Figure 5.**
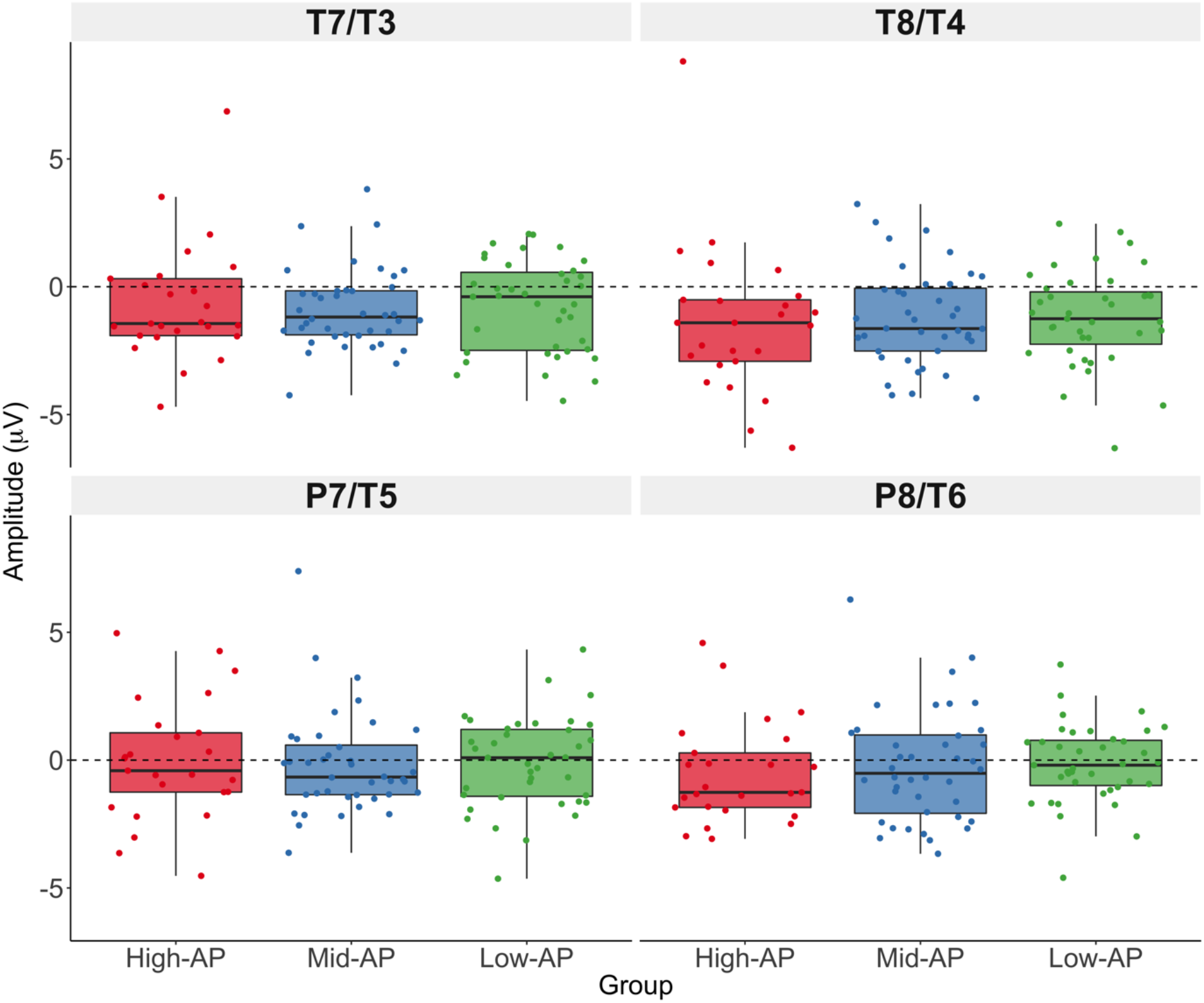
Lack of AP negativity elicitation. We could not replicate the main finding of the original study — the elicitation of an AP negativity in High-AP musicians in the Listening condition at electrode P7/T5 (bottom-left panel). Furthermore, there were no group differences at other temporal electrodes during the AP negativity-specific time window (bottom-right panel and upper panel). This is consistent with the original study, in which the AP negativity showed a high topographic specificity and was not elicited at electrodes T7/T3, T8/T4, and P8/T6.

#### AP Negativity – Control Analysis

To ensure that the diverging results regarding the AP negativity were not caused by the slightly different EEG preprocessing in this study, we reran the complete analysis using a preprocessing pipeline which closely reproduced the preprocessing steps undertaken in the original study. To this end, we re-referenced the raw EEG data to the linked mastoids, before correcting ocular artifacts using independent component analysis. Then, we segmented and baseline-corrected the data. Segments containing artifacts were rejected using a threshold of ±100 µV, and the clean segments were averaged to compute ERPs. Finally, we low-pass filtered the ERPs at 30 Hz (48 dB/octave). The statistical analysis of the AP negativity yielded virtually identical results; the one-way ANOVA at P7/T5 did not reveal a main effect of Group (F(2,101) = 0.006, p = 0.99, BF_01_ = 10.93, η^2^_G_ < 0.001). Thus, the results can be considered robust across different preprocessing pipelines.

## Discussion

The primary goal of this study was to evaluate the presence of an AP negativity in High-AP musicians, an important finding in AP research originally reported by Itoh et al. (2005). Using a large sample size, we failed to replicate the elicitation of this AP-specific ERP component during tone listening. Bayesian inference in the form of Bayes factors indicated moderate evidence (BF_01_ > 3) in favor of the null hypothesis that no AP negativity was elicited in High-AP musicians (or in any of the other groups). We also found strong evidence in favor of the null hypothesis of no group differences in the AP negativity (BF_01_ > 10). The evaluation of the ERP correlates of RP yielded mixed results. We were able to replicate the group differences in the P3b at electrode Pz. Low-AP musicians showed a higher P3b than Mid-AP musicians, but there were no differences between Mid-AP and High-AP musicians. Thus, different from the original study, there was no gradual increase in P3b amplitude with lower levels of AP. Also, in contrast to the original study, the P3b was elicited in all groups including High-AP musicians. An identical pattern of results was found at Fz. With regard to the ppSW, we replicated the group differences at electrode Pz. Again, Low-AP musicians showed a higher ppSW than Mid-AP musicians, but there were no differences between Mid-AP and High-AP musicians. Unlike in the original study, all groups elicited a ppSW including High-AP musicians. Finally, we were not able to replicate the group differences in the fnSW. Bayes factors indicated moderate evidence for the null hypothesis of no fnSW group differences at electrode location Fcz (BF_01_ > 3).

Before turning to the detailed discussion of the findings of this replication study, it is crucial that we delineate the differences between this study and the original study. Several of these differences are effectively improvements we implemented to mitigate methodological issues of the original study. First and foremost, a critical difference between the two studies is that in Itoh et al.’s sample, the groups did not only differ in terms of AP proficiency, but also in musical experience. In fact, there was a statistically significant difference in years of musical training between Mid/High-AP musicians and Low-AP musicians; this difference makes it difficult to disentangle the differential effects of AP and musical experience as it has repeatedly been shown that musical training influences auditory evoked potential components (Fujioka et al., 2006; Pantev et al., 1998; Schneider et al., 2002). In this replication study, the groups were matched with regard to musical experience (see Table 1). In a similar vein, none of the subjects in Itoh et al.’s sample were professional musicians, whereas our sample predominantly consisted of professional musicians and music students; this is also reflected in the substantial difference in years of musical training between Itoh et al.’s sample (mean of all AP groups = 11.33 years) and our sample (mean = 19.82 years). Since the groups in this replication study were matched for years of musical training, the influence of this characteristic was controlled for and thus cannot account for the differential results. As the original study was conducted in Japan and this study was conducted in Switzerland, the two samples also differ with regard to their ethnicity. Furthermore, the sample of the original study predominantly consisted of female subjects, whereas the sex distribution in our sample was balanced (49% females). Nevertheless, in general, it is implicitly assumed that the conclusions drawn from the sample of a study are generalizable to the whole population of interest. In the original study, the authors thus made conclusions about the whole population of AP possessors and assumed that these conclusions would also hold for AP possessors irrespective of sex and different genetic and cultural background.

Moving on from the characteristics of the two samples, there are also differences regarding the statistical analyses that must be mentioned. In ERP research, the definition of the analyzed time window for a component is an essential analysis step. Itoh et al. defined the time windows based on a visual comparison of the group-specific waveforms. In these waveforms, the authors identified the components that differed between the groups and subsequently performed the statistical analysis on the time windows in which these differing components appeared. This procedure is an example for what Luck and Gaspelin (2017) call “multiple implicit comparisons”, where the visual comparison of the group-specific waveforms correspond to a large number of statistical tests without an adjustment of the significance level. At the same time, this procedure makes the statistical analysis contingent on the results because the choice of the time window is not independent of the results. This is known as circular analysis which results in invalid inference and inflated effect sizes (Kriegeskorte et al., 2009; Vul et al., 2009). In cases of circular analysis, replications are crucial because they can provide a completely independent assessment of the extent to which the non-independent choice of the time windows had an influence on the results (Luck and Gaspelin, 2017). In this study, we thus used the time windows as defined in the original study. Another issue of the statistical analyses employed by Itoh et al. is the assumption of a difference between groups when one group showed a significant effect and another group showed a nonsignificant effect (Gelman and Stern, 2006; Nieuwenhuis et al., 2011). We thus performed statistical tests to directly compare the groups, in addition to the tests performed in the original study (more details are given in the Methods section).

As noted in the introduction, many studies cite Itoh et al.’s finding of an AP negativity at the left-sided electrode P7/T5 as evidence for the importance of the left planum temporale in AP, not least because this interpretation was also given in the original study. This rationale is not compatible with the current standard of knowledge regarding the neuronal sources of scalp ERPs. Most importantly, ERPs at a particular electrode, for example the P7/T5 which is placed over the left posterior temporal cortex, are not simply generated by the neural activity in the patch of cortex directly underlying this electrode. On the contrary, the relationship between ERPs and its neuronal sources is extremely complex; even highly sophisticated algorithms are only able to provide rough estimations of these sources (Pascual-Marqui et al., 2009). As Itoh et al. did not perform source estimation, the finding of an AP negativity in the original study should not be interpreted as evidence for the role of the left planum temporale in AP.

After having outlined the differences between the two studies, we now focus on the implications the results of this replication study have. How well does the non-elicitation of an early occurring AP-specific ERP component fit with evidence from previous EEG studies on AP? More than 25 years ago, Tervaniemi et al. (1993) investigated the mismatch negativity (MMN) in AP musicians. The MMN is an early ERP component elicited between 50 and 250 ms after stimulus onset and occurs in auditory oddball paradigms where a frequently presented standard stimulus is contrasted with an infrequently presented deviant stimulus. In their seminal study, Tervaniemi et al. (1993) found no differences in the MMN amplitude between musicians with and without AP that were matched in musical training. This finding was replicated in two independent studies (Rogenmoser et al., 2015). In a further study on the magnetic equivalent of the N1 component, Pantev et al. (1998) found differences between musicians and non-musicians, but again no differences between musicians with and without AP. The lack of differences in early auditory evoked potentials is also consistent with a more recent EEG study, in which there were no differences in the N1-P2 complex between AP musicians and Non-AP musicians (Elmer et al., 2013). Taken together, the non-elicitation of an early AP-specific component is compatible with a number of previous findings; a lack of early differences in auditory evoked potentials between musicians with and without AP as in this replication study seems to be the rule rather than the exception.

With regard to the ERP correlates of RP, the results of this replication study are in line with previous findings from EEG studies on AP. The reduced P3b component in subjects with higher levels of AP is not only consistent with Itoh et al.’s findings; it is also consistent with a number of EEG studies demonstrating a reduced or even absent P3b component in AP possessors during auditory oddball paradigms (Hantz et al., 1992; Klein et al., 1984; Wayman et al., 1992). A larger P3b amplitude is often associated with larger working memory demands (Polich, 2007). Labeling tones using RP necessarily involves working memory processes because the currently perceived tone needs to be compared to previously perceived tones; from a (small) set of tones, the name of the tone can then be inferred by comparison after the whole set of tones was perceived at least once. Further evidence for the role of working memory in RP processing comes from functional imaging studies showing stronger activation in musicians without AP in brain regions such as the posterior part of the inferior frontal gyrus during the labeling of tones (Zatorre et al., 1998). The interpretation of lower ppSW amplitudes in subjects with higher levels of AP is less straightforward and the role of the ppSW in RP is unclear. Itoh et al. interpreted the ppSW component as a reflection of the process to associate the pitch stored in working memory and its label. At the same time, the authors provided little evidence to support this interpretation. As the ppSW component is highly dependent on the P3b (see waveforms in Figure 3), it is also possible that it reflected further working memory processing (García-Larrea and Cézanne-Bert, 1998). Future studies should try to elucidate the potential roles of the ppSW in RP. Lastly, in this replication study, the fnSW was not different between High-AP, Mid-AP, and Low-AP musicians. One reason for this non-replication could be that in the original study, this component was only elicited in the groups which had lower levels of musical training (Low-AP and Untrained). It is thus plausible that the elicitation of an fnSW only occurs in individuals with little to no musical training and that it might be unrelated to both RP and AP processing.

Since the very beginning of scientific AP research, there has been a discussion about the status of musicians who in tone-naming tests perform significantly better than chance-level but worse than highly proficient AP musicians (Bachem, 1955). These musicians have been termed *Quasi-AP* (Bachem, 1937) or *AP-2* (Baharloo et al., 1998; Loui et al., 2012, 2010). In both the original study and this replication study, these musicians most likely make up a large proportion of Mid-AP musicians. However, the categorization of subjects to subgroups according to certain cutoffs in tone-naming proficiency is completely arbitrary. Furthermore, it is unclear whether Mid-AP musicians are actually RP musicians using a strategy (e.g., using the memorized pitch of the tuning standard A4 in combination with RP processing to infer the pitch of a tone) or whether they are AP musicians performing badly in the tone-naming test for some unobservable reason (e.g., because they are lacking motivation). In that sense, it is crucial to additionally consult the self-report of the subjects as this can help distinguish AP from RP musicians. In this study, we repeatedly observed a lack of differences between High-AP and Mid-AP musicians. Additionally, Low-AP musicians performed only marginally above chance level in tone-naming and can thus be characterized as RP musicians (cf. Materials and Methods). Taken together, the findings of this replication study suggest that a categorization into more than two groups, i.e. AP and RP musicians, may not be necessary. This notion of two separate populations for AP and RP musicians is also supported by large-scale behavioral studies employing tone-naming tests (Athos et al., 2007; Wengenroth et al., 2014).

A limitation of this study concerns the sample size. We invested considerable time and resources to recruit a large number of subjects (n = 104). However, due to the categorization of the subjects into three groups, the statistical power for some tests were slightly compromised. For example, even though descriptively there were differences at electrode Pz between High-AP and Low-AP musicians in both the P3b and the ppSW, these differences did not reach statistical significance and Bayes factors indicated inconclusive evidence (i.e. no hypothesis should be preferred). Inconclusive evidence, in turn, signifies the need for more subjects. As the recruitment of even more AP musicians is an extremely difficult task due to the rarity of the phenomenon, collaborations between multiple research groups should be considered to distribute the workload and foster the recruitment of even larger samples. A further limitation is that this study was not preregistered. Preregistration refers to the pre-specification of the hypotheses, the sample, the experimental procedure, and the statistical analyses of a study. These pre-specifications are then published on a website before the study is conducted. In general, preregistration protects against publication bias and analytic flexibility (Munafò et al., 2017). As we implemented some modifications to the pitch-naming task and, more importantly, to the statistical analyses, we could have preregistered these modifications to further increase the confidence in our findings.

To conclude, in this replication study using a large sample size, we were not able to replicate the elicitation of an AP negativity in High-AP musicians during tone listening, a landmark finding in neuroscientific AP research. We partially replicated the larger elicitation of the P3b and the ppSW, ERP correlates of RP, in musicians with lower levels of AP. As neither an isolated original study nor an isolated replication study can provide a final verdict on the veracity of a finding, more replication studies are imperative to further elucidate the neural underpinnings of AP.

## Acknowledgements

We thank our research interns Anna Speckert, Fabian Demuth, Florence Bernays, Joëlle Albrecht, Kathrin Baur, Laura Keller, Melek Haçan, Nicole Hedinger, Pascal Misala, Petra Meier, Sarah Appenzeller, Tenzin Dotschung, Valerie Hungerbühler, Vanessa Vallesi, and Vivienne Kunz for their invaluable help in data acquisition and research administration. Furthermore, we thank Anja Burkhard and Christian Brauchli for their support in the context of the larger absolute pitch project. Finally, we thank Carina Klein, Stefan Elmer, and all other members of the Auditory Research Group Zurich (ARGZ) for their valuable comments on the experimental procedure.

